# Still standing: persistence traits capture belowground plant functions beyond resource exploration and acquisition

**DOI:** 10.64898/2026.05.06.723249

**Authors:** Shersingh Joseph Tumber-Dávila, Karl Andraczek, Daniel C. Laughlin, Helge Bruelheide, Aline B. Bombo, Ying Fan, Alessandra Fidelis, Grégoire T. Freschet, Lena Hartmann, Justus Hennecke, Cody Coyotee Howard, Saheed O. Jimoh, Jitka Klimešová, Liesje Mommer, Tsumbedzo Ramalevha, Frances Siebert, Alexandra Weigelt, Joana Bergmann

**Author notes:** Corresponding Author: Shersingh Joseph Tumber-Dávila.

## Abstract

Belowground plant trait research has predominantly focused on trade-offs in fine root traits via the root economics space. Yet, this fine root framework captures only a fraction of the functional strategies plants employ beneath the soil surface. Here, we broaden the perspective on belowground plant functioning by integrating traits related to root system extent, clonality and bud banks, using data from the new UNDERPLOT database. This integration links measurable traits to key belowground functions: resource acquisition, spatial exploration, and persistence. Our analysis shows that the fine root economics space explains less than 5% of the variation in traits related to root system extent, clonality, and bud banks. Instead, an expanded trait analysis reveals three significant dimensions, explaining 62% of total trait variation. The third dimension, represents an independent, persistence-related gradient, not captured by existing root economics frameworks. We propose that understanding belowground plant strategies requires embracing additional functional gradients. The strategy of persistence, in particular, varies significantly across growth forms and is a critical dimension of plant response to resource limitation and stress, becoming increasingly important as global change shifts disturbance regimes.

## Introduction

Spectrums of plant form and function have been studied for decades (Grime, 1979), and advances have been made with the advent of global plant trait databases (Reich, 2014; Díaz *et al*., 2016; Klimešová *et al*., 2017a; Kattge *et al*., 2020; Guerrero-Ramírez *et al*., 2021; Iversen *et al*., 2021). Still, studies that seek universal patterns typically use a limited set of traits determined by existing functional knowledge but also their availability in databases. Belowground plant parts have received less attention than aboveground parts, and research on the relationships between traits remains fragmentary (Freschet & Roumet, 2017; Laliberté, 2017; Weigelt *et al*., 2021; Carmona & Beccari, 2025). Most knowledge on belowground plant functioning has been restricted to a few traits, often studied in isolation. For example, research on fine root traits rarely considers other forms of belowground trait variation such as those relating to clonality or bud bank size, which prevents us from achieving an integrative understanding of plant form and function (Freschet *et al*., 2021; Chelli *et al*., 2024; Liu *et al*., 2025; Matthus *et al*., 2025).

Two major drivers of belowground plant trait evolution are resource limitation and disturbance (Grime, 1979; Laughlin, 2023; Liu *et al*., 2025). Integrating traits that reflect key functional strategies to cope with these drivers is essential for understanding the relationships and trade-offs among belowground organs from an evolutionary perspective. Pioneering works have introduced the leaf economics spectrum (Wright *et al*., 2004) and the global spectrum of plant form and function that linked “fast–slow” strategies of resource acquisition across leaves, stems, and roots (Reich, 2014; Díaz *et al*., 2016) while recent studies have sought to unify above- and belowground strategies by merging the global spectrum of plant form and function with root system trait data (Weigelt *et al*., 2021; Carmona *et al*., 2021; Beccari & Carmona, 2024). Various independent efforts have already linked meaningful proxy traits to the following functions: i. nutrient acquisition strategies reflected by the collaboration and the conservation gradient of the fine-root economics space (Bergmann *et al*., 2020; Matthus *et al*., 2025), ii. water acquisition strategies reflected by rooting depth and extent (Schenk & Jackson, 2002; Fan *et al*., 2017; Tumber-Dávila *et al*., 2022; Bachofen *et al*., 2024), and iii. multiplication and vegetative regeneration after disturbance reflected by budbank and clonal traits (Klimešová & De Bello, 2009; Ott *et al*., 2019; Klimešová *et al*., 2025). However, belowground functional traits—from root system architecture and size to clonality and bud bank traits—are often studied in isolation, limiting conceptual integration across trait dimensions.

Most individual studies usually focus on a set of related traits rather than exploring a range of traits representative of alternative strategies to satisfy certain plant functions. For example, studying plant water uptake dynamics requires capturing aspects of plant strategies related to whole root system size (Tumber-Dávila *et al*., 2022) but also localized space exploration at the fine-root scale (Freschet *et al*., 2021). Furthermore, clonal growth organs and belowground buds might provide alternative strategies for horizontal space occupancy (Klimešová *et al*., 2018; Chelli *et al*., 2024) on top of their functional role related to vegetative reproduction especially across disturbance-prone ecosystems in which these strategies are most prevalent (Klimešová *et al*., 2017b; Pausas *et al*., 2018).

Here, we aim at broadening our perspective on the multidimensionality of belowground plant trait space that matters for plant functioning across its lifespan. To start from first-order principles, we organize belowground plant functions and propose traits that capture these functions. Using ecological knowledge on the different belowground trait categories (fine-root traits, root system size, and clonality/bud banks) as well as respective data at a global scale from the UNDERPLOT database (Bruelheide *et al*., in prep), we develop and test a conceptual framework that links traits to belowground plant functions and broader functional gradients (Fig. **1**). Based on existing knowledge, we hypothesize three distinct aspects of plant functioning to be covered by different belowground traits. These aspects (and their related functions in parentheses) are: (i) **spatial exploration** (resource exploration & anchorage), (ii) **resource acquisition**, and (iii) **persistence** (multiplication & regeneration). These functions are relevant at different scales of biological organization, i.e. that of the whole individual and that of particular organs, such as fine roots.

**Figure 1.**
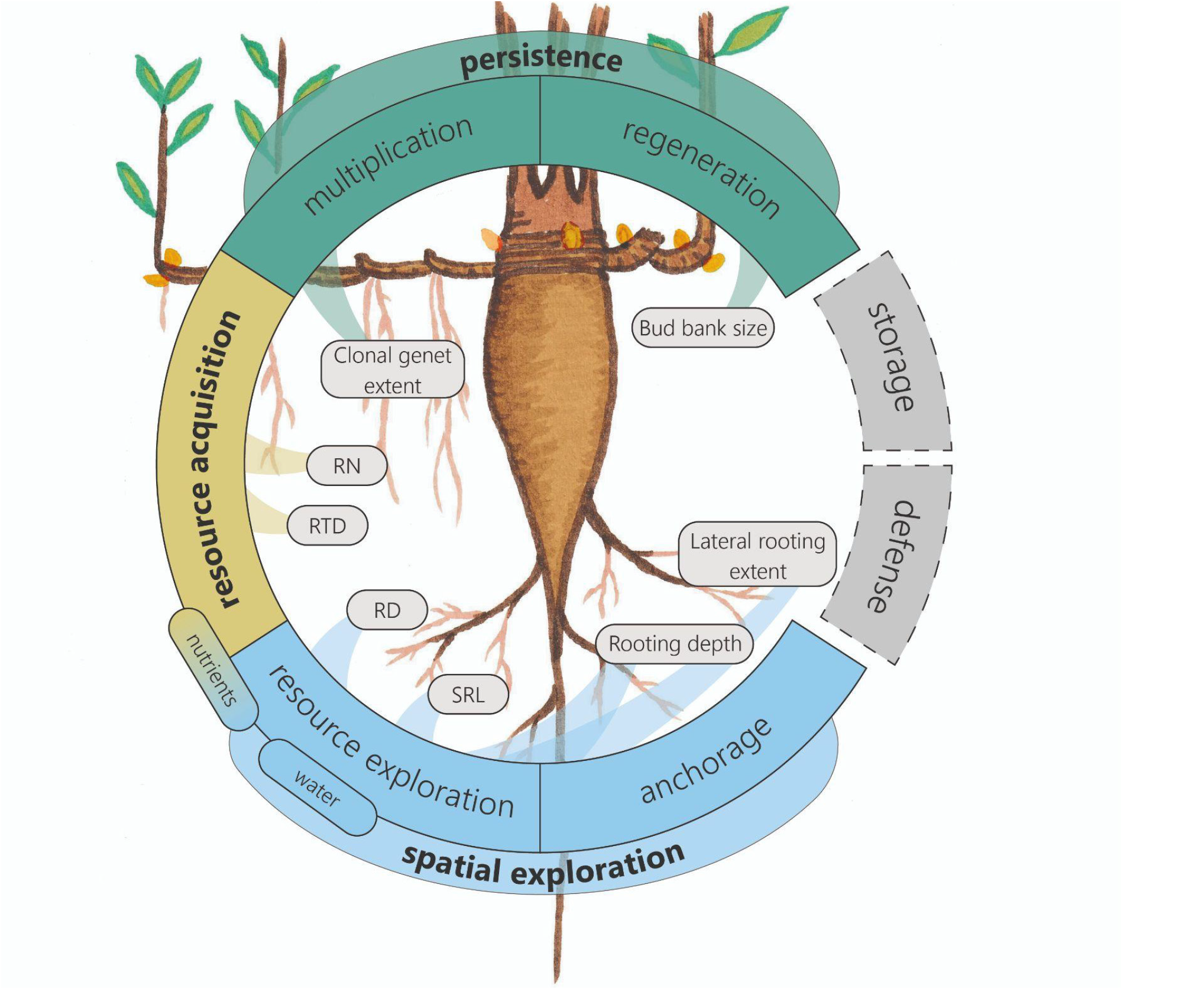
Conceptual linkages between traits and plant functioning leads to three dimensions of variation in the belowground functional space. The three colors (blue, yellow, green) reflect the three functional gradients hypothesized and identified in the data analysis (i.e., spatial exploration, resource acquisition, and persistence) and include either one or two related functions. Functions in grey (i.e. storage, defense) could not be incorporated into the analysis due to a lack of data, and so the conceptual model does not cover all potential functions that could govern belowground trait coordination. Traits in the center of the circle (described in Table **S1**) are a non-exhaustive set of measurable proxies mainly for one, but in some cases for two functions (e.g. lateral rooting extent and rooting depth serving as proxies for resource exploration and anchorage). The fine root trait abbreviations are: SRL - specific root length; RD - average root diameter; RTD - root tissue density; RN - root nitrogen concentration.

### Spatial exploration

Spatial exploration of the soil is critical for the acquisition of water and nutrients but also for anchorage in the soil. Resource exploration can be maximized at the whole-root system level by increasing maximum rooting depth (RDepth) and/or lateral rooting extent (LRExtent; Schenk & Jackson, 2002; Fan *et al*., 2017; Tumber-Dávila *et al*., 2022). Additionally, fine root morphology is crucial for spatial soil exploration for water and nutrients (Freschet *et al*., 2021; Kwatcho Kengdo *et al*., 2022). Maximizing absorptive surface area by increasing specific root length (SRL) or forming large diameter fine roots (RD-fine root diameter) to collaborate with mycorrhizal partners are two alternative strategies along the “collaboration gradient” which is equivalent to the *resource exploration* function at the scale of single fine roots in our conceptual model (Fig. **1**; Bauhus & Messier, 1999; Ostonen *et al*., 2007; Bergmann *et al*., 2020; Freschet *et al*., 2021).

Anchorage in the soil is mainly provided by root system traits (rooting depth and lateral rooting extent), which represent the maximum amount of space occupied by the plant belowground (Ennos, 1989; Mickovski *et al*., 2007; Tumber-Dávila *et al*., 2022). However, the horizontal space occupied by roots (i.e. lateral rooting extent) has been demonstrated to be highly related to the size of plants aboveground, the mechanics of which provide more anchorage support than rooting depth alone (Ennos, 1993; Tumber-Dávila *et al*., 2022). Rooting depth, especially in seasonally dry environments or places with deep yet accessible water tables is tied more directly to water uptake (Fan *et al*., 2017; Stocker *et al*., 2023; Bachofen *et al*., 2024; Mailloux *et al*., 2025). An increase in specific root length can further support anchorage, especially in the upper soil layers (Ennos, 1993), strengthening the functional link between resource exploration and anchorage in a common dimension of spatial exploration. Alternatively, for clonal plants, horizontal space occupancy belowground is primarily achieved via clonal spread and belowground stems (Klimešová *et al*., 2018; Klimešová, 2025).

### Resource Acquisition

Within the concept of the root economics space (RES), the conservation gradient shows variation between (i) fast nutrient acquisition by high metabolic activity reflected by high root nitrogen content (RN) in cheap but short-lived roots, and (ii) long lasting nutrient conservation in high tissue density roots (RTD-root tissue density) that are costly to construct (Bergmann *et al*., 2020). This gradient of variation between different strategies of resource acquisition has been shown to align with the aboveground fast-slow economics spectrum (Reich, 2014; Weigelt *et al*., 2021; Matthus *et al*., 2025) and captures temporal variation in carbon investment for resource acquisition. Apart from resource acquisition, plant functional variation along the conservation gradient has consequences for plant interactions with various soil biota (Lin *et al*., 2024; Neyret *et al*., 2024; Barry *et al*., 2025). Species with fast root traits (high RN) are generally considered to experience stronger negative plant-soil feedbacks (Semchenko *et al*., 2022; Spitzer *et al*., 2022; Delory *et al*., 2024). Furthermore, species with fast root traits affect soil carbon cycling at the community level (Bardgett, 2017; Barry *et al*., 2025), in particular by promoting increases in aboveground production through rapid resource acquisition (Da *et al*., 2023).

### Persistence

Vegetative regeneration and multiplication serve the higher order function of genet persistence that resembles a novel dimension in the belowground trait space. The belowground is generally a safe place for plants to survive environmental disturbances as the majority of disturbances hardly scratch the soil surface. Plants therefore position their regenerative organs belowground filled with storage carbohydrates and covered in buds, allowing them to persist at the site. The reliance on vegetative regeneration of once established individuals, or on regeneration of a new generation from seeds, depends on plant strategy of how to cope with a particular disturbance regime. A framework of Belowground Persistence Types (BPTs) was proposed (Klimešová et al. 2025) to summarize how different plant strategies cope with severe disturbance. The types are based on woodiness, clonality and resprouting ability and range in ascending degree of individual persistence from 1) herbaceous seeders, 2) herbaceous non-clonal resprouters, 3) herbaceous clonal resprouters, 4) woody seeders, 5) woody non-clonal resprouters to 6) woody clonal resprouters (Methods Note **1.1**). The categories of seeders (1 and 4) rely on seed reproduction and do not invest into belowground storage and bud bearing organs, while other plants rely on clonal organs and vegetative resprouting (BPT 2,3,5, and 6), regenerating when aboveground plant parts are destroyed by disturbances (i.e., fire, herbivory, flooding, or seasonal adversity; (Klimešová & Klimeš, 2007; Lubbe *et al*., 2021; Schnablová *et al*., 2025). Therefore, we use bud bank size (i.e. the number of buds per shoot) to represent resprouting capacity as a proxy for *regeneration*. While only one of many understudied bud bank traits, bud bank size is the most widely measured and is strongly linked to plant architecture and adaptation to disturbance regimes or plant stressors (Fidelis *et al*., 2014; Pausas *et al*., 2018; Ott *et al*., 2019; Bombo *et al*., 2024; Te *et al*., 2025).

Furthermore, we propose clonal genet extent (i.e. distance between the youngest and oldest connected ramet of a clonal plant) as an aggregate trait to quantify the horizontal dimension of a clonal plant, i.e. how much space the genet can occupy. For this we combined two traits: lateral spread, the distance achieved via clonal spread in one year, and the longevity of rhizomes responsible for the lateral spread. Therefore, clonal genet extent is linked to belowground persistence but is not a perfect metric for overall genet size or age, which are difficult to measure for clonal plants (Vonlanthen *et al*., 2010; de Witte & Stöcklin, 2010; Klimešová *et al*., 2026). Multiplication and spatial exploration via clonal growth also diminishes the role of the root system in resource exploration, thereby shifting the role of roots primarily towards resource acquisition (Chelli *et al*., 2024; Klimešová & Herben, 2024). Therefore, persistence strategies belowground can be only understood by jointly assessing the roles that bud banks, clonality and the root system play to promote carbohydrate storage, protection from disturbance, and spatial exploration (Klimešová & Herben, 2023, 2024).

#### Additional functions and conceptual considerations

We acknowledge that storage of resources and chemical defenses of belowground organs (gray boxes in Fig. **1**) are two key functions not directly informed by our set of traits (Araki *et al*., 2020; Lubbe *et al*., 2021), but we note that bud bank size is partly serving the function of storage in addition to regeneration. However, we have limited data and coverage for traits that serve as suitable proxies for the functions of storage and chemical defense (Hennecke *et al*., 2023; Bassi *et al*., 2024). Belowground functions may be considered along with suitable proxy traits in the future, for example, storage organ size seems to be a promising surrogate to storage function (Bartušková *et al*., 2022; Harris *et al*., 2025; Schnablová *et al*., 2025). There are additional aspects of belowground plant functioning and carbon storage not incorporated in this framework, such as exudation, rhizodeposition, and fungal symbionts, all of which are directly tied to plant functions such as resource acquisition and storage (Matthus *et al*., 2025; Wen *et al*., 2026).

While available traits being an approximation will never perfectly resemble a function, further data coverage of specialized traits will be needed to solidify our conceptual thinking. Though our conceptual framework considers the identified traits being linked to only one functional gradient, many traits may serve additional functions than the primary ones hypothesized herein.

#### Evaluation of the conceptual framework

We evaluated our conceptual framework with a newly synthesized UNDERPLOT database (Methods Note **1**; Fig. **S1**; Bruelheide *et al*., in prep) which has data from over 10,577 vascular plant species spanning 21 functional traits to determine whether the selected traits form distinct functional gradients of belowground plant functions. We began by testing whether the dimensions explained by the root economics space capture the total belowground trait space, or if the addition of single traits representative of root system size and clonality/bud banks lead to additional dimensions that better capture broader belowground plant functioning, as hypothesized in our conceptual framework (Fig. **1**). Next, drawing in all of the identified traits from all three belowground trait categories (fine-root traits, root system size, and clonality/bud banks), we determined how many dimensions were needed to capture belowground plant trait functioning, and if these new dimensions align with the functional gradients described in the conceptual framework.

#### The fine root economics space alone inadequately captures belowground trait variation

The root economics space concept has been widely developed and used to explain trade-offs in belowground plant functioning based on the coordination of plant traits organized into the collaboration (DIY-outsourcing; SRL-RD) and conservation (fast-slow; RN-RTD) gradients (Bergmann *et al*., 2020; Matthus *et al*., 2025). Here we assess to what extent additional traits related to belowground plant size (rooting depth and lateral rooting extent), clonality (clonal genet extent), and resprouting ability (bud bank size) can broaden our understanding of belowground plant functioning, by capturing independent dimensions beyond those characterized by the RES.

Using trait-specific regression models appropriate to the distribution of each trait (Methods Note **2.1**), we explore the association of the belowground traits mentioned above with the first two principal component axes of the RES-based PCA (Fig. **S2**,**S3a &** Table **S2**). We find that each of the additional traits vary systematically along the RES axes (*P* < 0.05), however the RES axes explain < 1% of the variance in rooting depth, lateral rooting extent, clonal genet extent, and bud bank size. The low proportion of variance explained suggests that RES traits alone cannot fully capture belowground functional variation, indicating a higher dimensionality of trait space.

#### Clonality and bud banks add a third dimension to the belowground trait space

Given that RES traits alone cannot fully capture the dimensionality of belowground plant functioning (Fig. **S2**; Matthus *et al*., 2025), we conducted a PCA with all traits from Fig. **1**, comprising the trait groups that reflect the three basic aspects of belowground plant form (fine roots, root system size, and clonality/bud banks; Fig. **2**). A third significant dimension emerged, with three principal components (with eigenvalues > 1) explaining 63% of the variation in belowground plant traits (Fig. **2** **&** Table **1**). Aligned with the RES, principal components (PC) 1 (26.5% of variance explained) and PC2 (18.5%) were largely governed by fine root and root system trait trade-offs (Fig. **2a**); however, the third significant dimension was driven largely by clonal genet extent, with PC3 accounting for 17.5% of the total variance explained (Fig. **2b** **&** Table **1**). Therefore, clonality, and in part bud bank size, make up a third independent dimension of belowground plant trait coordination that aligns with the persistence gradient of our conceptual framework, corresponding to the functions of multiplication and regeneration (Fig. **1**,**2b**). The permutation-based tests show that all traits aligned significantly with the first three principal components (*p* < 0.001; Methods Note **S2**), indicating that these axes capture major patterns of variation across belowground traits.

**Figure 2.**
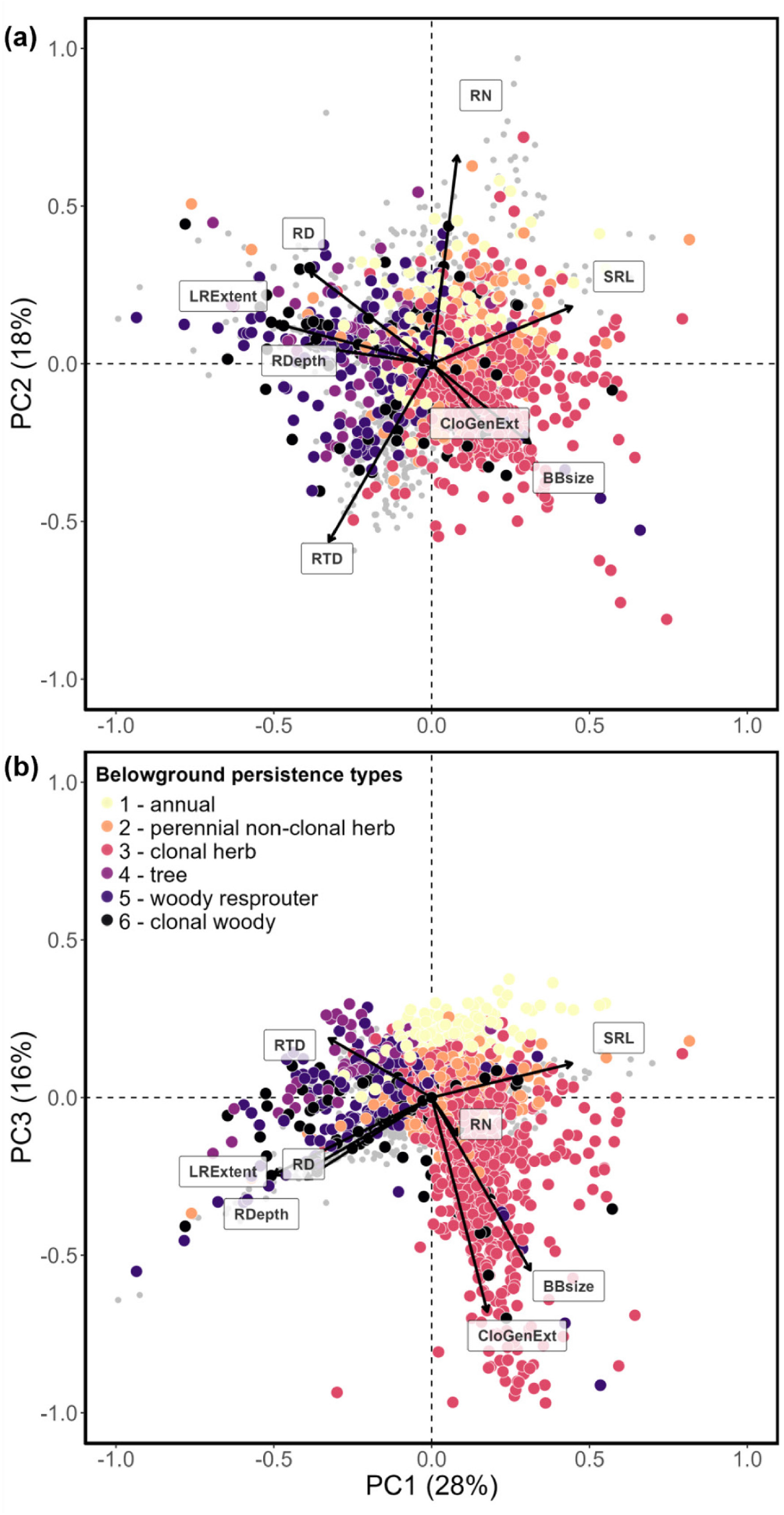
PCA on all traits reflecting the three main root functions for the three main significant PCs with points representing species colored by BPT. Trait data were obtained from the UNDERPLOT database (Bruelheide et al., in prep.). (a) PC1 and PC2 are driven largely by the root economics space and root system size, whereas (b) shows PC3 is driven largely by clonal and resprouting traits. The gray points are species that could not be assigned to a BPT due to missing trait information. For visualization only, species with missing RES trait values were projected into PCA space by replacing missing values with zeros prior to projection, which assigns mean trait values and biases their positions toward the center of the ordination. These species are shown in grey to indicate species coverage in our dataset and are excluded from all statistical analyses. The PC loadings for each trait are shown in Table **1**.

**Table 1:** Eigenvalues, variance explained, and loadings for the PCA in Fig. **2**.

|  | PC1 | PC2 | PC3 | PC4 |
| --- | --- | --- | --- | --- |
| Eigenvalues | 2.12 | 1.48 | 1.4 | 0.9 |
| Proportion of variance | 26.5 | 18.5 | 17.5 | 11.3 |
| Cumulative proportion | 26.5 | 45 | 62.5 | 73.8 |
| Root diameter | -0.39 | 0.24 | -0.27 | -0.59 |
| Specific root length | 0.45 | 0.20 | 0.04 | 0.47 |
| Root nitrogen content | 0.10 | 0.63 | -0.21 | 0.16 |
| Root tissue density | -0.34 | -0.53 | 0.23 | 0.26 |
| Maximum rooting depth | -0.45 | 0.24 | 0.25 | 0.35 |
| Lateral rooting extent | -0.47 | 0.15 | -0.15 | 0.28 |
| Clonal genet extent | -0.22 | 0.36 | 0.59 | -0.03 |
| Bud bank size | 0.23 | 0.09 | 0.63 | -0.37 |

Clonal genet extent and bud bank size appear to occupy additional dimensions of belowground plant functioning that is critical for maintaining genet persistence for certain species. While clonal genet extent load strongly on PC3, bud bank size has the highest proportion of unexplained variance across the first three principal components (Fig. **S3b**), which may reflect that higher dimensionality is needed to fully incorporate belowground functioning. Increase in bud bank size is the primary mechanism to enhance the ability to resprout after disturbance, and the unexplained variance in bud bank size could reflect additional functions such as the plants ability to store carbohydrates in order to respond to disturbances (Klimešová & Klimeš, 2007; Ott *et al*., 2019). Resprouting ability, along with defense and storage, are poorly understood mechanisms of belowground plant functioning, and all are functions related to resilience to disturbances. An increase in clonal genet extent promotes the probability of successful vegetative propagation near the mother plant (Klimešová *et al*., 2018; Ott *et al*., in press). However, both clonal genet extent and bud bank size are positively associated along PC3 (Fig **2b** **&** Table **1**), suggesting that the investment in multiplication is usually accompanied by an investment in regeneration with a result of increasing genet persistence. This indicates that clonality, which is restricted to about half of examined species, is nevertheless so important for shaping plant strategies that its absence or presence and its rate are affecting whole trait space.

#### Root economics is driven by trade-offs in spatial exploration and resource acquisition

The first axis of the integrated belowground PCA depicts a trade-off between specific root length on one side and fine root diameter, rooting depth, and lateral rooting extent on the other (Fig. **2a**). This partly conflicts with findings by Weigelt et al. (2021) and Beccari and Carmona (2024) who found that rooting depth loaded on separate and potentially multiple axes. Further complicating the dimensionality of the belowground trait space, trait-trait correlations (Fig. **S1**) reveal that few traits are actually varying independently, which leads to traits loading on more than their main axis of variation (Fig. **S3**).

At the root system scale, based on trait-plant functioning relationships, we argue that deep roots allow for access to deep soil water while lateral roots maximize the uptake of rain-fed water from upper soil layers, and both provide anchorage in the soil. As such, both rooting depth and lateral extent provide a proxy for whole-system spatial exploration and utilization of soil. Specific root length and fine root diameter—as fine-root traits—represent different strategies for small-scale spatial soil exploration for water and nutrient uptake (Freschet *et al*., 2021; Da *et al*., 2023; Barry *et al*., 2025). Our results suggest that whole-system spatial exploration of the soil is higher in species with wider fine root diameters that outsource small-scale exploration to mycorrhizal hyphae than in species with longer specific root length indicative of a “do-it-yourself” strategy (Bergmann *et al*., 2020). Furthermore, our PCA revealed resource acquisition—depicted by root tissue density and root nitrogen content—loads on PC2, hence being independent of spatial exploration and the collaboration gradient as has been proposed previously through the root economics space (Weigelt *et al*., 2021; Matthus *et al*., 2025).

#### Time to dig: future directions

Our analysis demonstrates that belowground plant trait variation requires at least three dimensions to capture the functions of spatial exploration, resource acquisition, and persistence. Through our conceptual framework, we further propose that functions such as *storage* and *defense* are important, but we so-far lack the trait data coverage and trait-function links to directly inform this. Therefore, this multidimensional framework represents an initial step toward a more comprehensive understanding of belowground plant functioning, and we acknowledge several limitations that point toward productive avenues for future research.

### Data limitations and the need for expanded trait coverage

While the UNDERPLOT database has advanced the synthesis of trait data across fine roots, root system size, and clonality/bud banks, substantial gaps remain (Bruelheide *et al*., in prep). Most clonal trait data originated from the CLO-PLA database and a limited number of additional sources that target underrepresented regions (Klimešová *et al*., 2017a; Pausas *et al*., 2018; Bruelheide *et al*., in prep); however, this still results in restricted overlap between root traits and clonal traits across species. Despite these limitations, our analysis revealed striking patterns that support the conceptual framework, suggesting that expanded data collection efforts will likely strengthen rather than contradict these findings.

Critically, two key belowground functions, resource storage and defense, could not be adequately incorporated into our framework due to insufficient trait data (Fig. **1**). While we acknowledge that bud bank size partially serves storage functions alongside regeneration, traits that more directly capture carbohydrate storage capacity (such as nonstructural carbohydrate content in storage organs) and belowground physical and chemical defense compounds (e.g. lignin-to-N content, root cellulose, phytochemical diversity) remain poorly represented in global databases (Bassi *et al*., 2024; Jimoh *et al*., 2024). Future data collection initiatives should prioritize traits as proxies for these underrepresented functions to enable a more holistic understanding of belowground plant strategies.

### Integrating above- and belowground perspectives

The belowground trait space has long been treated as a collection of disconnected trait dimensions studied in isolation, limiting our capacity to understand the integrated strategies plants employ beneath the soil surface (Iversen, 2014; Bardgett, 2017; Gallagher *et al*., 2020; Matthus *et al*., 2025). A logical next step involves linking this belowground conceptual framework with aboveground functional traits—especially for persistence which have not been represented in research linking above- and below-ground trait trade-offs—which have historically captured a broader range of functions beyond resource uptake. Of particular interest is understanding how vegetative persistence strategies (*multiplication* and *regeneration* via belowground organs) relate to genetic persistence through generative reproduction (Klimešová, 2025; Klimešová *et al*., 2025). This integration would bridge the persistent disconnect between above- and below-ground plant ecology while providing insights into the trade-offs plants face in allocating resources to different persistence strategies.

Furthermore, while recent efforts, such as the findings presented here, have begun merging the plant economics spectrum with root system trait data, the incorporation of clonality and resprouting traits into these unified frameworks remains nascent (Carmona & Beccari, 2025; Ott *et al*., in press). Our identification of persistence as an independent third dimension suggests that future syntheses must move beyond the traditional fast-slow and collaboration-conservation gradients to fully capture plant functional diversity, especially for disturbance-driven systems.

### A new research agenda

We propose that advancing our understanding of belowground plant functioning requires: (i) expanded trait data collection, particularly for functions with no clear trait proxies, such as storage and defense-related traits; (ii) increased overlap in species and spatiotemporal coverage across plant trait categories that reflect basic belowground functions— i.e., spatial exploration, resource acquisition, and persistence; (iii) explicit testing of trait-plant function relationships through manipulative experiments; (iv) integration with aboveground trait frameworks; and (v) examination of how environmental gradients and disturbance regimes shape the multidimensional belowground trait space. As global change accelerates and novel disturbance regimes emerge, understanding the belowground strategies that enable plants to persist, recover, and spread will become increasingly critical for predicting vegetation dynamics and ecosystem resilience. Only through such coordinated efforts can we advance and achieve the integrative understanding of plant form and function that has long remained elusive for the hidden half of plants.

## Acknowledgements

This paper is a joint effort of the working group sUnderfoot kindly supported by sDiv, the Synthesis Centre for Biodiversity Sciences at the German Centre for Integrative Biodiversity Research (iDiv) Halle-Jena-Leipzig, funded by the German Research Foundation (FZT 118, 202548816). S.J.T.D would like to acknowledge partial support from the LTER Postdoctoral Fellowship through Harvard Forest NSF Funded LTER Program (NSF-DEB-LTER 1832210), and additional input on the manuscript from the TD Lab and K. Harpenau. J.K. was supported by Praemium Academiae awarded by Academy of Sciences of the Czech Republic. A.F. was supported by Fundação de Amparo à Pesquisa do Estado de São Paulo (FAPESP 2023/16620-0), and Conselho Nacional Desenvolvimento Científico e Tecnológico (CNPq 312689/2021-7). GTF was supported by the Laboratoire d’Excellence TULIP (ANR-10-LABX-0041).

## Data Availability Statement

The data included in UNDERPLOT dataset are described in Bruelheide *et al*. (submitted) and are openly available under the terms specified by CC BY 4.0 (https://doi.org/10.25829/idiv.3610-56acy9). The R code used for the analysis is included as supplemental file in Rmarkdown format: “Data Analysis Script-Still standing.Rmd”.

## Supporting Information (brief legends)

**Methods Note 1**. UNDERground PLant-Organ Trait database – UNDERPLOT

**Methods Note 1.1**. The use of Belowground Persistence Types (BPTs)

**Methods Note 2**. Principal component and statistical analyses

**Methods Note 2.1**. RES PCA

**Methods Note 2.2**. Integrated belowground PCA

**Table S1**. Trait and species data from the UNDERPLOT Database used in this study

**Figure S1**. Correlation matrix of observations

**Figure S2**. RES PCA

**Table S2**. Eigenvalues, variance explained, and loadings for PCA in Fig. S2

**Figure S3**. Relative importance of PCA axes for belowground traits

**Figure S4**. PCA across plant classifications

