## Supporting Information for "Still standing: persistence traits capture belowground plant functions beyond resource exploration and acquisition"

The following Supporting Information is available for this article:

### Supporting Information (brief legends)

**Methods Note 1.** UNDERground PLant- Organ Trait database - UNDERPLOT
**Methods Note 1.1.** The use of Belowground Persistence Types (BPTs)
**Methods Note 2.** Principal component and statistical analyses
**Methods Note 2.1.** RES PCA
**Methods Note 2.2.** Integrated belowground PCA
**Table S1.** Trait and species data from the UNDERPLOT Database used in this study
**Figure S1.** Correlation matrix of observations
**Figure S2.** RES PCA
**Table S2.** Eigenvalues, variance explained, and loadings for PCA in Fig. **S2**
**Figure S3.** Relative importance of PCA axes for belowground traits
**Figure S4.** PCA across plant classifications

#### **Methods S1.** UNDERground PLant- Organ Trait database - UNDERPLOT In order to synthesize trait data across the three dominant categories of belowground plant traits (fine root traits, root system size, and clonal and bad bank traits) we aggregated a global dataset of plant traits called UNDERPLOT (Bruelheide *et al.*, in prep). The goal of UNDERPLOT is to bridge the major disconnect between traits related to belowground plant functioning, with researchers often focusing on distinct components of belowground functioning in isolation. This open-access dataset provides an opportunity to consider different belowground trait categories simultaneously, at the global scale, thus potentially increasing our comprehensive understanding of the belowground trait space and trait-environment interactions. The compiled and harmonized dataset includes information from 10,592 vascular plant species and 21 plant functional traits, along with categorical and descriptive information about the growth forms of each of the plant species, and a taxonomic standardization that utilizes the sPlot4.1 backbone (Bruelheide *et al.*, 2019). While UNDERPLOT consists of a variety of databases from hundreds of original sources, a majority of the fine root trait data comes from GRooT (Guerrero-Ramírez *et al.*, 2021), root system size data from RSIP (Tumber-Dávila *et al.*, 2022), and clonal data from CLOPLA (Klimešová *et al.*, 2017a), with additional species information from TRY-v6 (Kattge *et al.*, 2020) and FungalRoot (Soudzilovskaia *et al.*, 2020). Tables **S1, S2** show the trait data from UNDERPLOT used in this manuscript.

**Table S1.** Trait and species data from the UNDERPLOT Database used in this study (adapted from *Bruelheide et al.*, 2026)

| Trait Name (Abbreviation) | Units | Description | Number of species | Mean | Q 25% | Median | Q 75% |
| --- | --- | --- | --- | --- | --- | --- | --- |
| **Fine Root Economic Space Traits** | | | | | | | |
| Root Nitrogen Content  (RN) | mg/g | Fine root nitrogen content by the mass | 1850 | 13.66 | 8.31 | 12.10 | 17.50 |
| Fine Root Diameter  (RD) | mm | Mean diameter of fine roots | 1844 | 0.42 | 0.23 | 0.35 | 0.51 |
| Root Tissue Density  (RTD) | g/cm^3^ | Mass of fine roots by the volume | 1629 | 0.30 | 0.16 | 0.26 | 0.39 |
| Specific Root Length  (SRL) | m/g | Length of fine roots by the dry mass | 2146 | 91.60 | 21.28 | 51.22 | 114.20 |
| **Root System Size Traits** | | | | | | | |
| Max Rooting Depth  (RDepth) | m | Maximum rooting depth | 2759 | 1.58 | 0.44 | 0.90 | 1.70 |
| Lateral Rooting Extent  (LRExtent) | m | Maximum horizontal distance of root system from the base of the stem (radius) | 1810 | 1.59 | 0.25 | 0.48 | 1.11 |
| **Clonality and Resprouting Traits** | | | | | | | |
| Bud Bank Size  (BBsize) | buds/shoot | Total number of buds for each plant | 3220 | 12.79 | 5.00 | 12.80 | 20.00 |
| Clonal Genet Extent  (CloGenExt) | cm*years | Distance of clonal spread in a year multiplied by the persistence, representing the maximum lateral distance occupied by clonal plants | 1919* | 24.35 | 2.00 | 13.00 | 35.20 |

**CloGenExt has 6,002 observations when including zeroes for non-clonal observations*

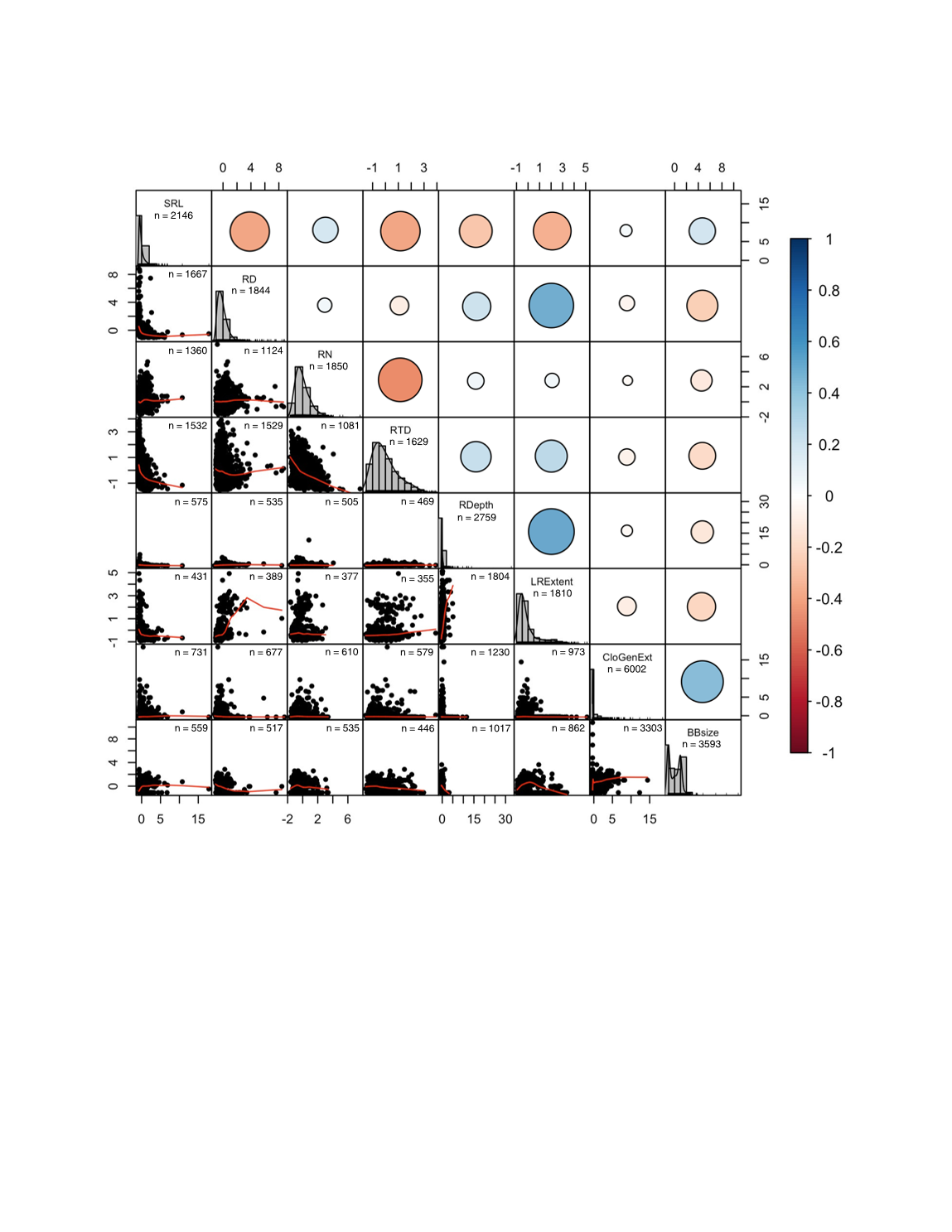

**Figure S1.** Correlation matrix of observations for specific root length (SRL), average root diameter (RD), root nitrogen concentration (RN), root tissue density (RTD), rooting depth (RDepth), lateral rooting extent (LRExtent), clonal genet extent (CloGenExt), and bud bank size (BBsize). Trait data were obtained from the UNDERPLOT database (Bruelheide e*t al.,* in prep.). Panels above the diagonal depict circles representing Pearson’s correlation coefficients for each trait-trait relationship, with size and color denoting magnitude and direction, respectively. Diagonal panels demonstrate histograms of trait values. Panels below the diagonal are trait-trait relationship scatter plots fit with red lowess curves. Values in the upper right and diagonal panels represent number of observations (n) for each trait pair. The traits used in this correlation matrix are described in Table **S1**.

**Methods S1.1.** The use of Belowground Persistence Types (BPTs)

To visualize how species with different life history strategies occupy the belowground trait space, we employed the Belowground Persistence Type (BPT) classification system introduced by Klimešová et al. (2025). We selected the BPT framework because it addresses gaps in previous life-form classifications by integrating both above- and belowground plant traits, particularly the role of bud banks and storage organs. The BPT framework categorizes terrestrial vascular plants into six functional types based on their adaptations to severe and recurrent disturbances, and overall plant genet persistence. This classification is structured around three binary traits: (i) woodiness, which reflects longevity and investment in plant structures, (ii) clonality, indicating whether a plant can regenerate through the production of genetically identical, physically independent units, and (iii) resprouting ability, which determines whether a plant can replace lost aboveground biomass following disturbance. The BPT categorization results in six plant functional types informed by both the above- and belowground characteristics of plants: (i) herbaceous seeders, (ii) herbaceous non-clonal resprouters, (iii) herbaceous clonal resprouters, (iv) woody seeders, (v) woody non-clonal resprouters, and (vi) woody clonal resprouters. This ordinal framework captures persistence and survival strategies of plants across disturbance regimes, ranging from short-lived herbaceous, annual seeders to long-lived, woody, clonal resprouters. BPT as demonstrated in Figs. **2**, **S2**, also conveniently integrates many of the binary traits of plant form (Fig. **S4**) into one ordinal system.

**Methods S2.** Principal component and statistical analyses

To identify to what extent a more holistic belowground trait space including fine root traits of the root economics space, as well as traits related to root system extent, bud banks and clonal organs, explain plant functional variation belowground, we conducted a stepwise ordination-based analysis. Specifically, we used Principal Component Analysis (PCA) using mean trait values per species based on complete empirical trait measurements. All trait data were log-transformed and standardized prior to analysis. All analyses were performed in R (version 4.4.0).

**Methods S2.1.** RES PCA
In a first step, we tested whether additional belowground traits represent independent axes of functional trait variation not captured by the RES traits. To do so, we computed a pairwise-complete correlation matrix based on the RES traits and performed a PCA using the `princomp` function (R Core Team, 2024) with a regularized covariance matrix, in which negative eigenvalues were replaced with small positive values. To assess whether additional belowground traits—namely maximum rooting depth, lateral rooting extent, bud bank size, and clonal genet extent—are associated with the axes of the RES or capture new dimensions of belowground plant functioning, we projected species into the RES space by multiplying their trait values with the PCA loadings of the RES ordination. The resulting PCA scores were centered (but not scaled) prior to downstream analyses.

We then quantified both the alignment of these traits with the RES axes and the proportion of trait variance explained by the RES. For rooting depth and lateral rooting extent, we used a vector-fitting approach analogous to the `envfit` function (Oksanen *et al.*, 2025). Each trait was regressed independently onto the first two PCA axes using linear models without intercepts. Trait significance was assessed by comparing the observed F-statistic to a null distribution generated from 999 permutations of trait values. Bud bank size and clonal genet extent were strongly zero-inflated, as these traits are only expressed in species that are clonal. These traits were therefore analysed using Tobit regression models with left censoring at zero. Model significance was assessed using permutation tests (999 permutations) based on the Wald statistic. For all traits, variance explained by PC1 and PC2 (representing the RES axes) was quantified using model R² for linear models, and McFadden’s pseudo-R² for Tobit models (Fig. **S3a**).

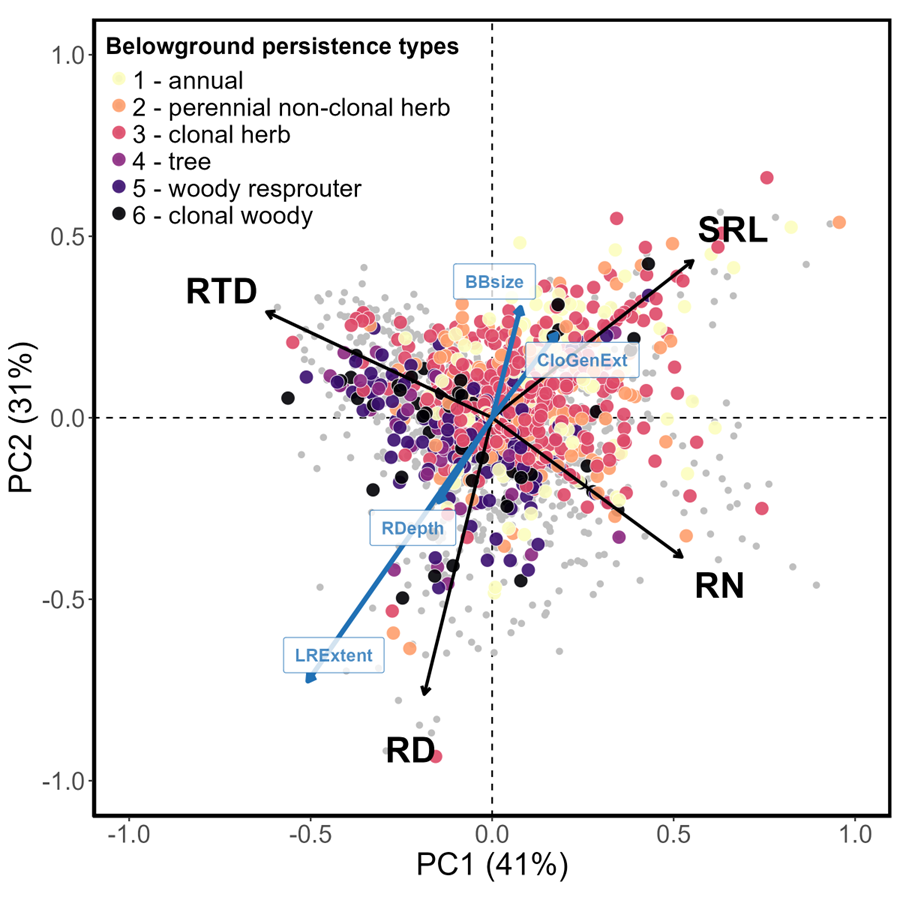

**Figure S2.** The RES PCA shows that the root economics space can be defined by two primary principal component axes that represent more than 70% of the variation in trait organizations of fine root traits (Black arrows). Trait data were obtained from the UNDERPLOT database (Bruelheide et al., in prep.). A post-hoc regression on axis scores was performed to see whether or not rooting depth (RDepth), lateral rooting extent (LRExtent), clonal genet extent (CloGenExt), and bud bank size (BBsize) fit within the RES (blue arrows). The colored points represent species trait averages among the RES color-coded by belowground persistence types (BPT; Klimešová *et al.*, 2025), with the gray points representing species that could not be assigned a BPT due to incomplete information. For visualization only, species with missing RES trait values were projected into PCA space by replacing missing values with zeros prior to projection, which assigns mean trait values and biases their positions toward the center of the ordination. These species are shown in grey to indicate species coverage in our dataset, but are excluded from all statistical analyses. The principal component loadings for each trait in the figure are shown in Table **S2**.

**Table S2**: Eigenvalues, variance explained, and loadings for PCA in Fig. **S2**.

|  | PC1 | PC2 | PC3 |
| --- | --- | --- | --- |
| Eigenvalues | 1.62 | 1.24 | 0.68 |
| Proportion of variance | 40.6 | 31.1 | 17 |
| Cumulative proportion | 40.6 | 71.7 | 88.7 |
| Root diameter | -0.19 | -0.76 | 0.46 |
| Specific root length | 0.55 | 0.43 | 0.42 |
| Root nitrogen content | 0.52 | -0.38 | -0.70 |
| Root tissue density | -0.62 | 0.29 | -0.35 |

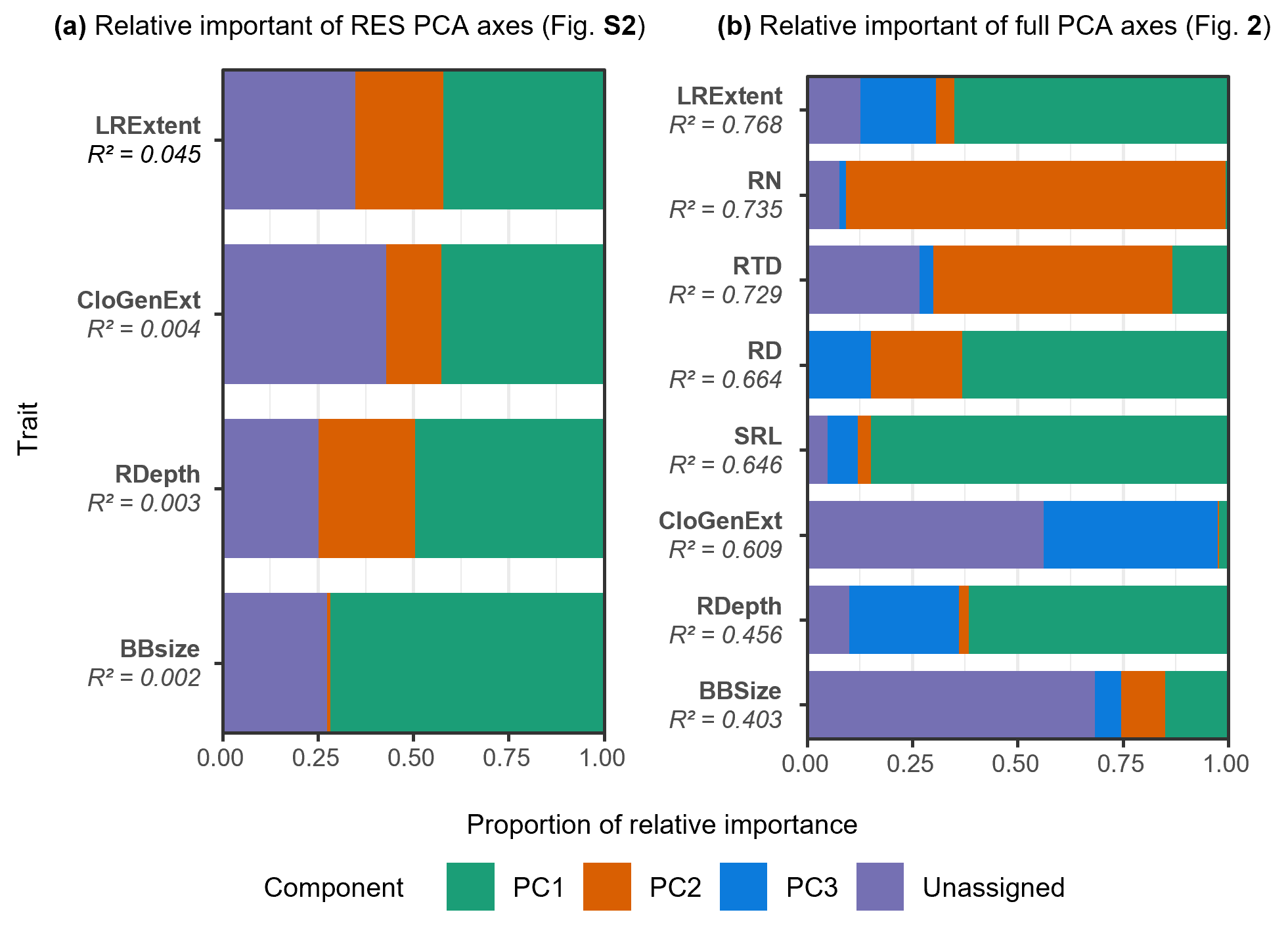

**Figure S3.** Relative importance of PCA axes for the RES PCA (**a**; Methods **2.1**) and for the full PCA (**b**; Methods **2.2**) for the main belowground traits. Traits are ordered based on the R^2^ of the linear fits for each model regressing the trait values against the PCA axes.

**Methods S2.2.** Integrated belowground PCA
In a second step, we aimed to identify the main axes of belowground functional trait variation across all traits jointly, including RES traits, maximum rooting depth, lateral root spread, bud bank size, and clonal genet extent. As described before, we computed a pairwise-complete correlation matrix. We then performed PCA on this regularized covariance matrix to derive the major axes of belowground trait variation. To interpret how individual traits aligned with these axes, we extracted PCA loadings for the first four components, and found three components to have eigenvalues greater than 1 (Table **2**). As done in M2.1, we quantified both the alignment of these traits and the proportion of trait variance explained by the first three principal components using a permutation-based regression analysis (Fig. **S3b**).

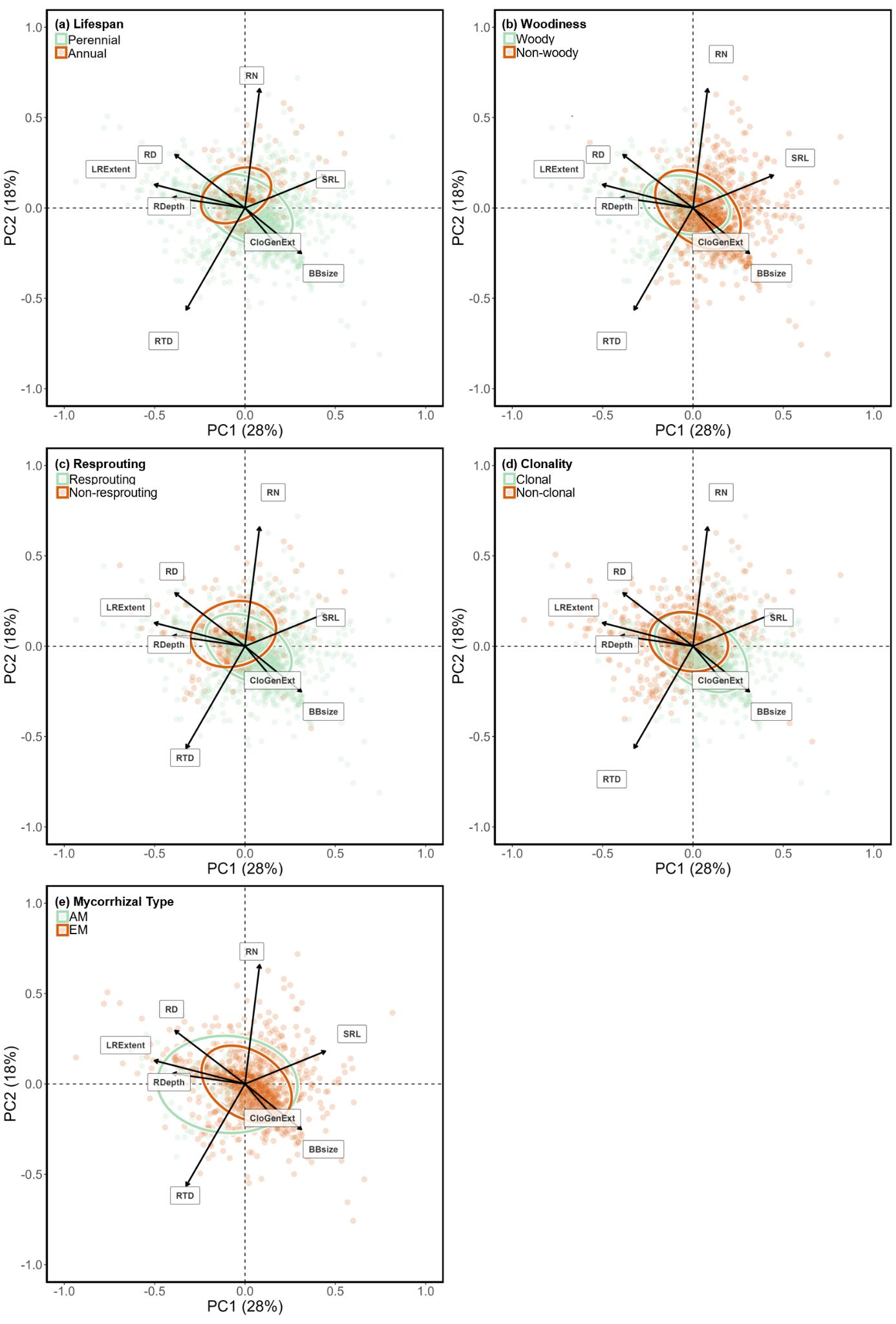

**Figure S4.** PCA on all traits colored by plant classifications of (a) Annual or Perennial, (b) Woodiness, (c) Resprouting, (d) Clonality, and (e) Mycorrhizal Type. This is the same PCA as in Fig. **2**, with only the classification/colors of the different species demarking the organization of species within those classes across the PCA.
